# *Streptococcus pneumoniae* augments circadian clock gene expression in zebrafish cells

**DOI:** 10.1101/2024.02.20.581283

**Authors:** Camila Morales Fénero, Raina E. Sacksteder, Jacqueline M. Kimmey

**Affiliations:** Department of Microbiology and Environmental Toxicology, University of California Santa Cruz

## Abstract

The circadian clock is a highly coordinated, cell-autonomous process that regulates the daily internal rhythms of biological organisms. Circadian clocks have been identified in various organisms, including mammals, invertebrates, cyanobacteria, and plants. The zebrafish (*Danio rerio*) is a practical model for studying the vertebrate circadian clock due to its small size, ease of manipulation, gene tractability, and direct light responsiveness. Several studies have revealed that bacterial, viral, and parasitic infections can impact the expression of circadian genes in mammals. While some evidence suggests that this may also be the case in zebrafish, no studies have investigated the direct effects of bacterial exposure on the zebrafish clock. Here, using zebrafish Z3 cells, we show that exposure to heat-killed *Streptococcus pneumoniae* (HK-Spn) can augment the expression of core repressive factors, including *per1b, per2, per3,* and *cry1a*. Further investigation demonstrated that HK-Spn induces the production of reactive oxygen species (ROS) in Z3 cells and that the addition of NAC, a ROS antioxidant, blocks Spn-mediated induction of *per2*, *cry1a*, and *per3*. Additionally, HK-Spn augmented the expression of *tefa* and *tefb,* an effect NAC suppressed. These results suggest the involvement of a ROS-dependent pathway in the augmentation of *per2*, *cry1a, and per3* by HK-Spn. Moreover, the activation of *tefa* and *tefb* by HK-Spn represents promising new targets for further investigation of the regulation of these genes.

## Introduction

Circadian rhythms are 24-hour oscillations that regulate different aspects of our physiology. In vertebrates, circadian rhythms are regulated by a molecular clock generated by a transcriptional/translational feedback loop (TTFL) consisting of activating and repressive regulatory complexes. The activation complex consists of the proteins Clock and Bmal, which drive the expression of genes containing E-box elements in their promoters. Clock/Bmal also drives the production of Per and Cry, which inhibits Clock/Bmal activity, leading to a negative feedback loop to terminate *Per* and *Cry* expression^1^. Importantly, environmental signals such as light can reset the rhythms of these genes in a process called entrainment, which enables organisms to anticipate and prepare for daily environmental changes^1–4^.

Zebrafish (*Danio rerio)* represent a fascinating model for studying key aspects of the vertebrate circadian system with characteristics like small size, easy genetic manipulation, and the ability of all the cells of their body and entire organs to autonomously entrain their rhythms by responding directly to light^5^. The molecular clock of the zebrafish has all essential components of the TTFL of vertebrates, and due to duplications in the teleost’s genome, some genes have two or more homolog copies^6–11^. In zebrafish, all cells are directly responsive to light, which induces *per2* and *cry1a* transcripts^12–22^. The current model of light regulation suggests that a photoreceptor detects light and activates the transcription factor thyrotrophic embryonic factor a (Tefa)^14^. Though this photoreceptor has not yet been identified, some candidates have been proposed including the teleost multiple tissue (*tmt*) opsin, flavin-containing oxidases, and Cry4^16,23–25^. Tefa is part of the proline- and acid-rich family of basic region leucine-zipper (PAR bZIP) transcription factors, which also includes albumin D-site-binding protein (DBP) and hepatic leukemia factor (HLF)^26^. Eleven PAR bZIP family genes have been identified in zebrafish, including *tefa* and its homolog gene, *tefb*^27^. *tefa* is induced by light and activates *per2* and *cry1a* through promoter elements called D-boxes^9,14,15,18,27,28^. However, while activation via D-boxes is sufficient to drive the expression of *cry1a*, the expression of *per2* requires both E-boxes and D-boxes, suggesting other factors participate in the light-induction of this gene^14,15,17^. Furthermore, overexpression of *tefa* in the dark is insufficient to induce *per2* expression^28^. Less is known about *tefb,* but a few studies have shown that *tefb* exhibits circadian rhythmicity and light regulation^15,27,28^. Knockdown of *tefb* alters light-induction of *per2*; however, as its putative DNA binding sites have not been determined, it is unknown if this results from direct promoter interactions with D-boxes^18,28^.

Reactive oxygen species (ROS) production has also been related to the light-induction pathway ^14–17,24^. ROS is a term used to describe a series of derivatives of molecular oxygen, which include free radicals such as superoxide radical anion, hydroxyl radical, and peroxyl radicals, as well as nonradicals like hydrogen peroxide (H_2_O_2_), singlet oxygen, and hypochlorous acid. ROS are generated by the electron transport chain in the mitochondria and NADPH oxidase (NOX), xanthine oxidase, and dual oxidase (DUOX) situated in different subcellular locations, including the plasma membrane, endoplasmic reticulum, mitochondria, and nuclear membrane^29–32^. Diverse factors, including metabolic cues, cytokines, and microbe recognition, can stimulate ROS generation^32,33^. Studies in zebrafish have shown that ROS generated in response to light activate MAP kinases to induce *per2* and *cry1a* expression through D-boxes^16,17,34^. Moreover, H_2_O_2_ is sufficient to induce the expression of *per2* and *cry1a*, which has been suggested to occur through the activation of D-boxes^16,17^.

As mentioned previously, the circadian clock constantly interacts with the environment, and a variety of inputs can change its expression. Different studies have shown that infection can disrupt clock gene expression in vertebrates, an effect that may alter disease outcome^35–38^. In rainbow trout (*Oncorhynchus mykiss*), infection with the ectoparasite *Argulus foliaceus* alters expression and rhythmicity of the clock genes, with differing effects depending on whether the fish were exposed to continuous light/dark (LD) cycles or constant light^39^. In zebrafish, RNA-seq analysis revealed that infection with *Pseudoloma neurophilia* downregulated the expression of multiple genes related to circadian rhythms in the fish’s brain, including *per1b* and *nr1d1*^40^. Studies in mammals also have shown evidence of changes in circadian gene expression during infection. For instance, human cells infected with the Hepatitis C virus (HCV) have decreased transcript and protein levels of PER2 and CRY2. When *PER2* was overexpressed, cells had lower levels of HCV replication and regained normal expression of interferon-stimulated genes, which are central components of anti-viral defenses^35^. Another study showed that infection with *Helicobacter pylori* up-regulated the expression of *BMAL1*, leading to an enhanced expression of TNF-α, aggravating the inflammatory response^36^. Finally, the clock genes *Bmal1*, *Per1*, and *Dbp* have decreased expression in infections with the eukaryotic parasites *Trypanosoma brucei* and *Plasmodium chabaudi* in mice^38^. These studies have shown the effects on circadian gene expression during active infection but raise the question of whether the observed effect is due to direct microbe exposure, inflammation, or damage caused by infection. Many studies have shown that cytokines can affect circadian rhythms, with pro-inflammatory molecules typically dampening circadian gene expression^41–44^. In zebrafish, a screen for small molecules that affect circadian rhythms using the transgenic line *per3:luc* also demonstrated amplitude-dampening effects upon exposure to pro-inflammatory compounds, while anti-inflammatory molecules had the opposite effect^45^. These reports indicate that inflammation indeed affects the circadian clock, yet no studies have investigated the direct effect of bacteria on circadian gene expression.

In this study, we utilized zebrafish Z3 cells to examine the impact of exposure to heat-killed *Streptococcus pneumoniae* (hereafter HK-Spn) on the expression of the repressive arm of the zebrafish clock: *per1b*, *per2*, *per3*, and *cry1a*. Our findings indicate that exposure to this bacteria increases the expression of *per2*, *per3*, and *cry1a* through a ROS-mediated pathway.

## Methods

### Z3 cell culture

Z3 cells, a fibroblast-like embryonic cell line from *Danio rerio*^12^, were maintained at room temperature in Z3 Media: Leibovitz L-15 medium (Gibco, 11415064) supplemented with 15% HyClone Fetal Bovine Serum (Cyvita, 16777-238), 2µM L-Glutamine (Sigma, G3126) and 1% Penicillin/Streptomycin (Sigma, P4333). Cells were seeded at 1×10^6^, 5×10^5^, or 5×10^4^ cells/well into 6-, 12- or 96-well plates, respectively, grown until confluent and entrained in 12:12 LD cycle for three days using low-intensity white light (LED, 400-700 nm, ∼500-lux). After entrainment, cells remained in the dark unless light was noted in the figure legend. A single media change was performed before entrainment.

### Light sources

Cells in light conditions were exposed to low-intensity (∼500-lux) or high-intensity (∼5000-lux) white light (LED, 400-700 nm) for 4 hours or 12 hours, as noted in the figure legend. Cells in dark conditions were kept in constant darkness and treated and harvested under very dim red light (LED, 600-750 nm, ∼0-2 lux), which does not induce *per2*.

### Heat-killed bacteria

Heat-killed *Streptococcus pneumoniae* (HK-Spn) was prepared by growing wild-type D39 in Todd-Hewitt Broth (Thermo Scientific, CM0189B) with 2% Yeast Extract (Sigma, Y1625) at 37°C with 5% CO_2_ until they reached an OD_600_ of 0.4. Colony-forming units were enumerated by serial dilution and plating on Columbia Blood Agar (Thermo Scientific, OXCM331B) with 5% Sheep’s Blood (Fisher Scientific, 50863755), and the remaining bacteria were then pelleted, resuspended in PBS, and heated to 100°C for 1 hour. HK-Spn was stored at −20 °C until use.

### Bacterial exposure and pharmacological treatments

PBS, HK-Spn, and 300µM H_2_O_2_ (Sigma, H1009) were applied at time zero. N-Acetyl-L-cysteine (NAC) (Sigma, A9165) was added 2 hours before the experiment started to a final concentration of 6 mM. Due to variation between batches of HK-Spn, the effective dose was determined empirically based on augmentation of *per2* and *cry1a* after 4 hours in light conditions. For all experiments shown here, the actual dose used was between 3.7×10^8^-2.25×10^9^ CFU/mL.

### RNA extraction and RT-qPCR

Cells were harvested in DNA/RNA Shield (Zymo, R110050) by scraping, flash frozen in liquid nitrogen, and stored at −80°C until extraction. The following products were used for qRT-PCR analysis: Quick-RNA Miniprep Kit (Zymo, R1055), iScript cDNA Synthesis (Bio-Rad, 1708891), PowerUp SYBR Green (Applied Biosystems, A25777), CFX Connect Real-Time thermocycler (Bio-Rad). Primers: *actb1* (Forward:5’-ATCTTCACTCCCCTTGTTCAC-3’; Reverse:5’-TCATCTCCAGCAAAACCGG-3’), *per1b* (Forward:5’-TGCGCGTAATGGAGAGTATATG-3’; Reverse:5’-CTTCGTTCAGTGGAGAGGTTC-3’), *per2* (Forward:5’-ACGAGGACAAGCCAGAGGAACG-3’; Reverse:5’-GCACTGGCTGGTGATGGAGA-3’), *per3* (Forward:5’-CAAGTACAAGCAAACAGCGAG-3’; Reverse:5’-ACTACCACAAAAGAGTCCGTG-3’), *cry1a* (Forward:5’-GGAGTGTGAACGCAGGAAG-3’; Reverse:5’-AAACCCCTTAAGACTGGCAG-3’), *tefa* (Forward:5’-TCACTCTGTTGCCTGTCATG-3’; Reverse:5’-CGTCTACCTCGATTTCATCTGG-3’) and *tefb* (Forward:5’-TGCTAATGATGCCTGGACAC-3’; Reverse:5’-TGCTAATGATGCCTGGACAC-3’). ΔΔCT was used to calculate relative mRNA expression^46^. Values were normalized to *actb1* and compared to time zero samples.

### Measurement of intracellular ROS

Prior to stimulation, cells were incubated in 10µM 2′,7′-Dichlorodihydrofluorescein diacetate (DCFH-DA) (Sigma, D6883) in Hanks Balanced Salt Solution (HBSS) (Gibco, 14025092) for 45 minutes. Cells were washed once to remove extracellular DCFH-DA and placed in plain HBSS, followed by treatment with HK-Spn, NAC, or PBS as indicated. All treatments were conducted under dim red light (∼0-2 lux). DCF was measured at 30-minute intervals at 490 nm excitation and 530 nm emission wavelengths in a Spectramax iD3 Multi-Mode Microplate Reader (Molecular Devices). Baseline normalization was performed by subtracting readings from wells without cells.

## Results

### Basal and light-induced expression of circadian genes on Z3 cells

In zebrafish, light directly induces the expression of *per2* and *cry1a* through promoter elements called D-boxes^14,15^. The acute production of these core repressive factors represents the principal mechanism by which zebrafish clocks entrains to new light cycles^13,18,19^. Here, we focused our research on the expression of these two light-driven genes, *per2* and *cry1a*, as well as two non-light-induced genes, *per3* and *per1b.* Because most studies have been implemented in PAC-2 cells, we first sought to verify whether the patterns of RNA expression of circadian genes are conserved in zebrafish Z3 cells, an embryonic fibroblasts-like cell line^14,15,17,18,27^. Z3 cells were subjected to 12:12 LD cycles of entrainment for three days, followed by an additional 12-hour dark period (during an expected light period). To verify that Z3 cells maintain circadian rhythms *in vitro* and retain the ability to respond to light, cells were subjected to a 12-hour light period (LDDD) or left in constant darkness (DDDD). We then analyzed the transcripts expression of *per1b, per2, per3,* and *cry1a* every 4 hours for 48 hours (Fig.1A).

**Figure 1.**
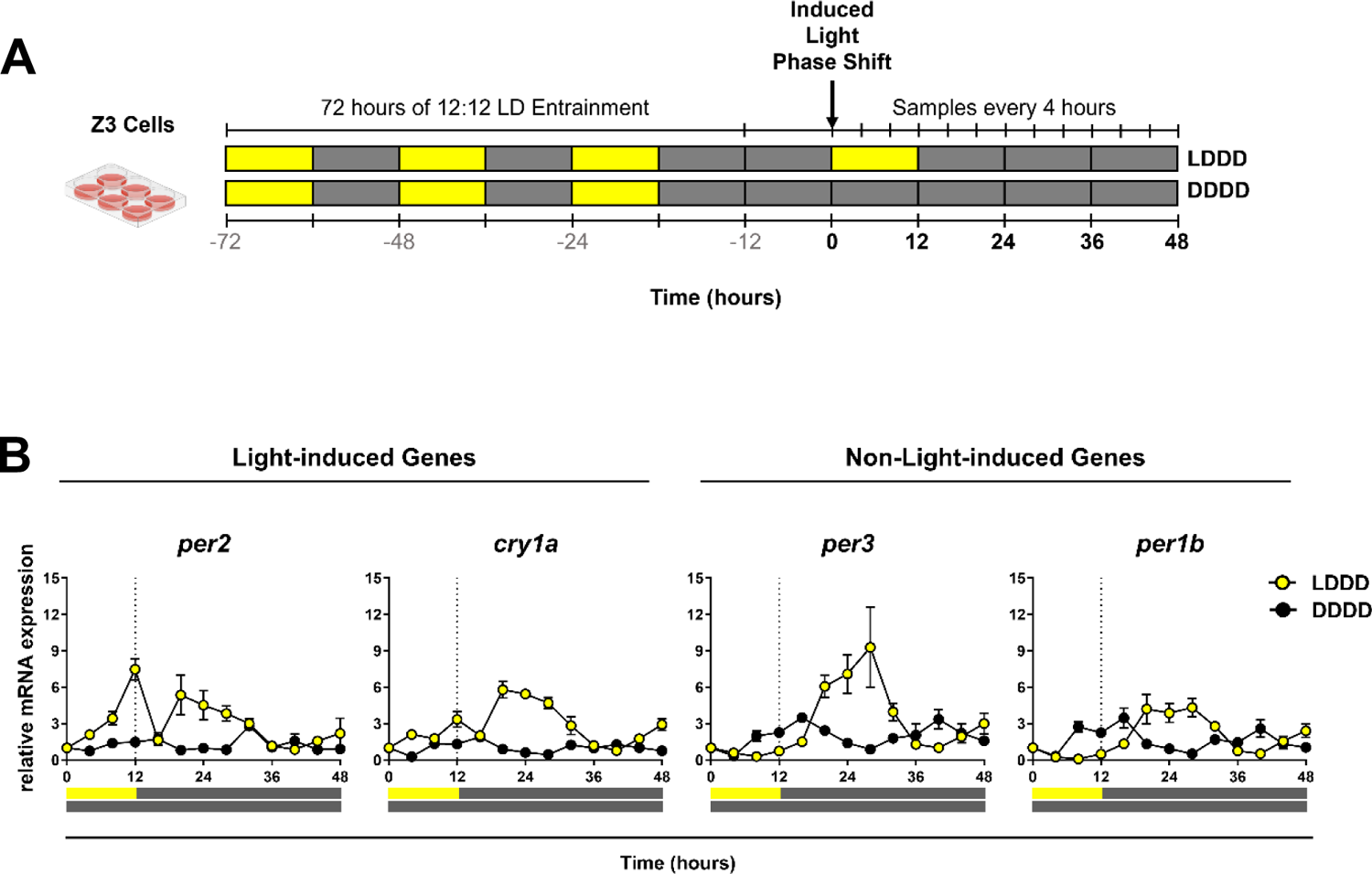
Light regulation of *per1b, per2, per3* and *cry1a* on Z3 cells. **(A)** Z3 cells were entrained for three light/dark (LD) cycles of 12:12 h, then phase-shifted, exposed to a 12-hour dark gap followed by either 12-hour light (∼500-lux) with subsequent darkness (LDDD) or to constant darkness (DDDD). RNA samples were taken every 4 hours for 48 hours. **(B)** mRNA expression of the light-induced genes *per2* and *cry1a* and the non-light-induced genes *per3* and *per1b* on LDDD (yellow circle) and DDDD (black circle) for 48 hours. Transcripts levels were measured against time zero. Yellow bar: light exposure. Grey bar: constant darkness.

As predicted, immediate increases in *per2* and *cry1a* occurred in response to light (0-12 hours) that is not seen in *per3* or *per1b* (Fig.1B). Light-induced expression occurs due to D-boxes in the promoters of *per2* and *cry1a*, and in line with previous reports, these genes are minimally expressed in the dark.^9,14,15,17,18,27^. Light did not induce the expression of *per1b* or *per3* (LDDD) but instead shifted their expression in comparison to constant darkness (DDDD), confirming that a 12-hour light period is sufficient to invert the circadian phase (Fig.1B). Prior reports in larvae and PAC-2 cells have shown *per1b* and *per3* are E-box regulated^21,47,48^. In addition to D-boxes, *per2* and *cry1a* also contain E-box sequences^14,15^, which likely explains why all four genes have similar expression patterns in DDDD (peak at 12 and 36 hrs) and a peak at 24hr in LDDD. These data validate the expression patterns of these genes previously shown in other cell lines.

### Heat-killed *Streptococcus pneumoniae* augment gene expression of the zebrafish clock’s repressive arm

Prior studies have demonstrated that viral, bacterial, and parasitic infections can affect the expression clock genes in mammals and teleost fish, and in some cases, this is dependent on the lighting parameters^35,36,38–40^. To determine if exposure to microbial ligands in the absence of a live infection directly affects the expression of clock genes, we exposed Z3 cells to heat-killed *Streptococcus pneumoniae* (HK-Spn), a gram-positive bacterium. Bacteria were added at time zero in the same lighting parameters as in Fig.1, and we monitored the effect of HK-Spn on *per2, cry1a, per3,* and *per1b* gene expression for 48 hours. As in Fig. 1, the expression of *per2* and *cry1a* was induced by light. Surprisingly, exposure to HK-Spn further amplified the light-induced expression of *per2* and *cry1a* (Fig.2A and S1). Moreover, the amplitude of *cry1a* expression was also augmented during the subsequent dark period (20-36hr). In contrast, there was no marked increase in amplitude in per3 and per1b expression in the LDDD condition.

**Figure 2.**
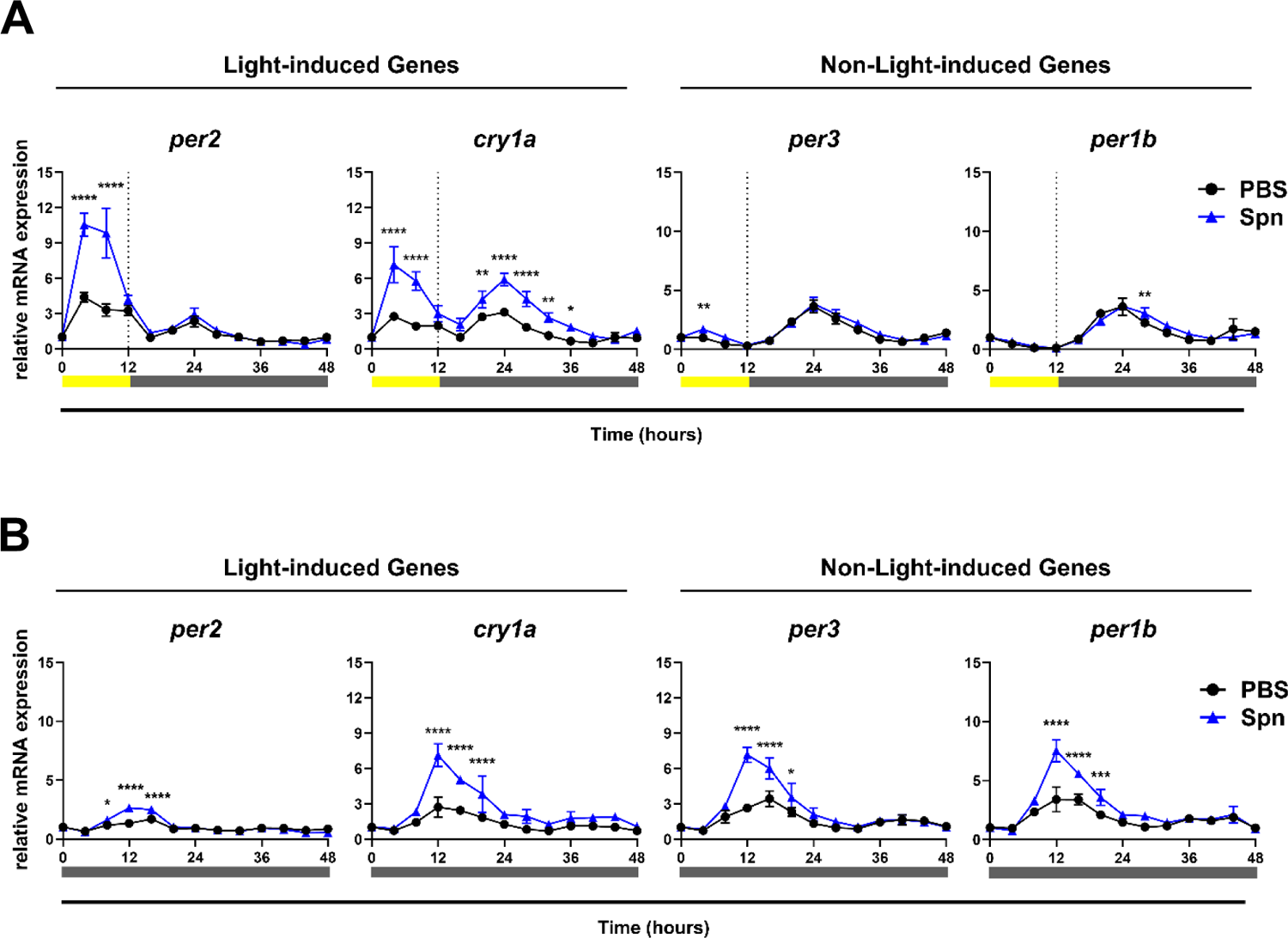
HK-Spn exposure augments the expression of *per2, cry1a, per3* and *per1b*. Z3 cells were entrained for three LD cycles of 12:12 h, followed by a 12-hour light pulse (LDDD) or in constant darkness (DDDD). Heat-killed *Streptococcus pneumoniae* (HK-Spn) was added at time 0, and samples were collected every 4 hours throughout 48 hours. **(A)** Transcript expression of *per2, cry1a*, *per3*, and *per1b* of PBS or HK-Spn*-*exposed cells after 12-hour light (500-lux) followed by constant darkness (LDDD). **(B)** Transcript expression of *per2, cry1a*, *per3,* and *per1b* of PBS or HK-Spn-exposed cells in constant darkness (DDDD). *p<0.05,** p<0.01,***p<0.001, ****p<0.0001, Two-way ANOVA with Šídák’s multiple comparisons test. Yellow bar: light period. Grey bar: dark period.

Based on these results, we suspected that exposure to HK-Spn may only affect the expression of light-induced genes (*per2* and *cry1a*) but not non-inducible ones (*per3* and *per1b*) due to interaction with D-boxes. We repeated these analyses in constant darkness to determine whether the effect depends on light. As expected, *per2* expression is modest under continuous darkness; however, a small but statistically significant augmentation occurred in response to HK-Spn (Fig.2B). HK-Spn substantially augmented the amplitude of *cry1a, per3* and *per1b* oscillations within the first 24 hours following exposure, a result confirmed in an independent 24-hour repeat of this experiment (Fig.S1). The effect is not mediated solely through D-boxes as *per3* and *per1b* both lack this regulatory region and suggest other motifs, such as E-boxes, might play a role in the response.

### HK-Spn acutely augments *per2, cry1a,* and *per3* in different light conditions

Given that *per1b, per2, per3,* and *cry1a* are part of a negative feedback loop that can affect their own expression, we decided to focus on the acute induction of genes within the first 4 hours of exposure. At this time point, the changes observed are most likely to stem from the direct effects of HK-Spn exposure. Because the intensity of light affects the expression of *per2* and *cry1a*^13,18^, and since the most prominent effect of HK-Spn exposure was seen in *per2* light-driven expression (∼10-fold increase), we next sought to determine whether HK-Spn would have the same effect at multiple light intensities. As in Fig.2B, HK-Spn is insufficient to induce expression in the dark of *per2* or *cry1a* at the 4-h timepoint. 500-lux light induces *per2* and *cry1a* expression (∼4.0-fold and ∼3.3-fold on average, respectively), which is further augmented in the presence of HK-Spn (∼10.2-fold and ∼6.5-fold on average, respectively) (Fig.3). Notably, *per2* and *cry1a* induction is strongly increased with high-intensity light (5000-lux) (∼13.6-fold and ∼4.8-fold, respectively), which is further magnified in the presence of HK-Spn (*per2* ∼29.8-fold, *cry1a* ∼9.5-fold).

**Figure 3.**
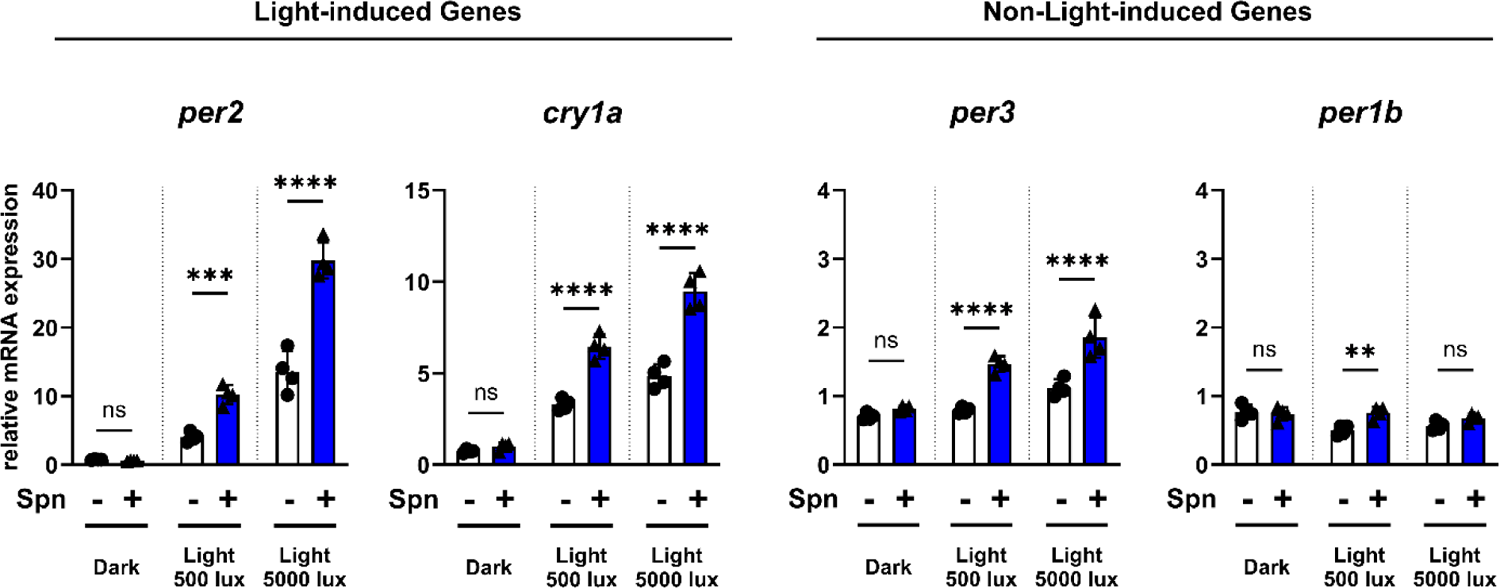
HK-Spn exposure acutely augments the light expression of *per2, cry1a,* and *per3*. Z3 cells were exposed to HK-Spn in dark and light conditions (500-lux and 5000-lux). RNA expression of *per2*, *cry1a*, *per3*, and *per1b* was measured after 4 hours. ** p<0.01,***p<0.001,****p<0.0001, Two-way ANOVA with Šídák’s multiple comparisons test between preselected groups.

In the case of *per3*, neither 500-lux light nor HK-Spn alone induces expression, but there was a small effect in combination (∼1.5-fold increase) (Fig. 3). High-intensity light modestly induced *per3* (∼1.1-fold induction, *p = 0.0032*) which was significantly augmented by the addition of HK-Spn (∼2-fold). As previously described, *per1b* is inhibited by light^11,21^, and in contrast to *per2, per3,* and *cry1a,* HK-Spn did not augment *per1b* expression. Based on this, we hypothesize that HK-Spn may only be able to augment genes being actively transcribed (i.e., *per2, per3,* and *cry1a* in the light) rather than being able to induce the expression of these genes independently.

### Exogenous H_2_O_2_ does not replicate light’s transcriptional conditions

Prior literature indicates that one mechanism by which light induces *per2* and *cry1a* is through ROS and that H_2_O_2_-mediated activation of D-boxes is sufficient to drive transcription of these genes in the dark^16,17^. We, therefore, wondered whether HK-Spn would also augment H_2_O_2_-mediated expression of *per2* and *cry1a* in the dark. Z3 cells were exposed to 300 µM H_2_O_2_ alone or in combination with HK-Spn in dark conditions. As expected, H_2_O_2_ induced robust expression of *per2* and *cry1a* at 4 hours post-stimulation; however, in contrast to light, there was no combinatorial effect of H_2_O_2_ and HK-Spn (Fig.4). H_2_O_2_ was insufficient to induce *per3* or *per1b,* consistent with previous reports indicating H_2_O_2_ activates D-boxes (absent in *per1b* and *per3*)^17^. However, in contrast to *per2* and *cry1a*, the combination of H_2_O_2_ and HK-Spn moderately augmented the expression of *per3* and *per1b*. We do not understand what distinguishes these genes to allow these differential responses. Regardless, it is clear that H_2_O_2_ does not recapitulate the same cellular conditions as light.

**Figure 4.**
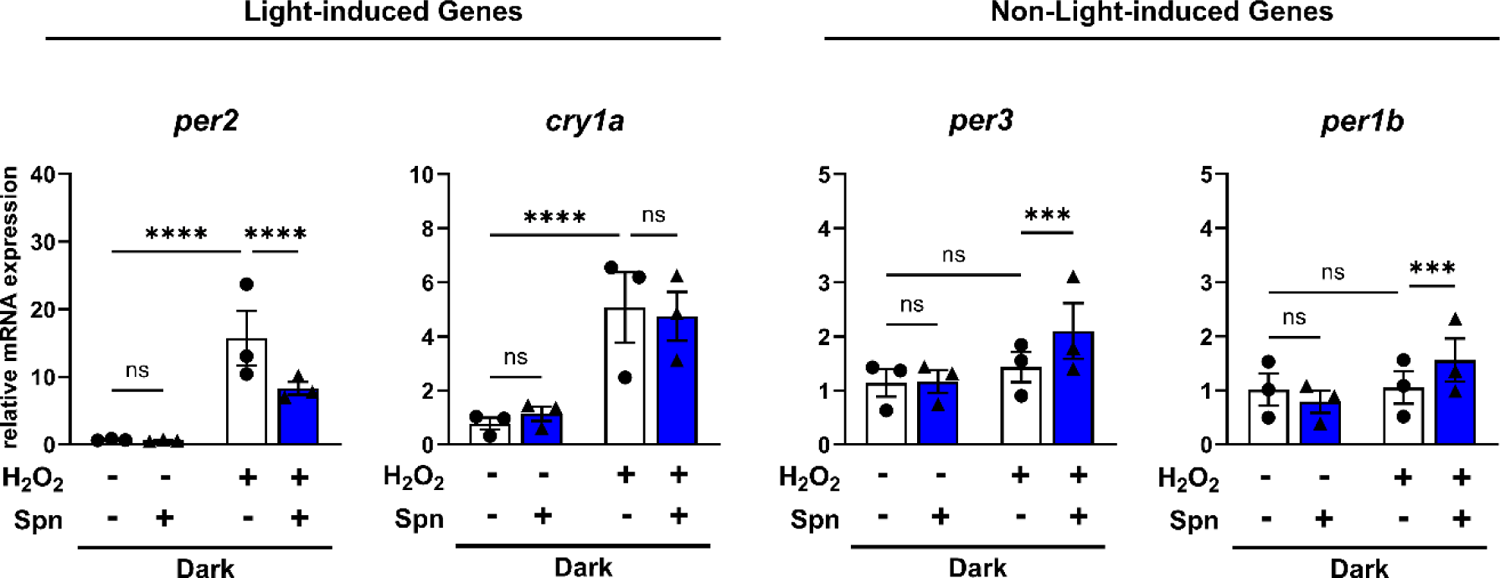
Hydrogen peroxide does not replicate the effects of light. Relative expression of *per2, cry1a, per3,* and *per1b* in Z3 cells incubated with 300 µM hydrogen peroxide (H_2_O_2_) for 4 in the dark.*** p<0.001, ****p<0.0001, Two-way ANOVA with Šídák’s multiple comparisons test between preselected groups.

### HK-Spn-induced ROS mediates *per2*, *cry1a,* and *per3* expression

Given that many cells generate ROS upon detecting microbial ligands^49,50^, we wondered if induction of intracellular ROS by HK-Spn might contribute to the augmentation of *per* and *cry* genes. DCFH-DA is a cell-permeable reagent that gets trapped in cells upon cleavage by intracellular esterases and fluoresces upon oxidation by intracellular ROS^51^. Z3 cells were incubated with DCFH-DA, washed, and exposed to HK-Spn. Figure 5A depicts independent kinetic graphs and quantified differences in the area under the curve of each condition.

**Figure 5.**
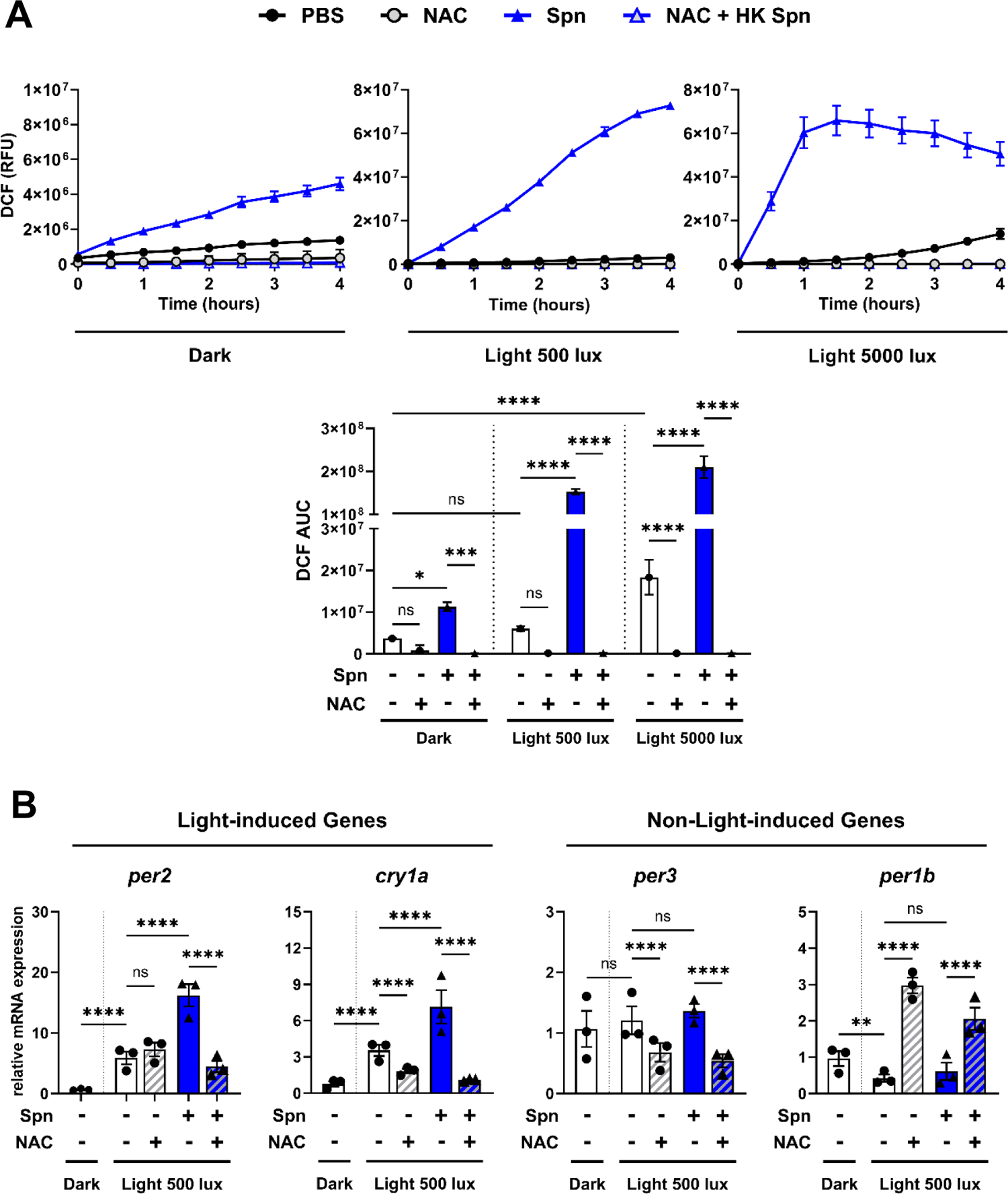
HK-Spn mediates *per2*, *cry1a,* and *per3* augmentation through ROS. **(A)** Relative fluorescent units (RFU) of Dichlorohydrofluoresin (DCF) were measured every 30 minutes for 4 hours on PBS or HK-Spn-treated Z3 cells in the dark, low-intensity light (∼500-lux), and high-intensity light (∼5000-lux). A 2-hour pre-treatment of 6 mM of N-acetyl cysteine (NAC) was used to suppress ROS production. The area under the curve (AUC) was measured and compared between groups and conditions. *p<0.05, ***p<0.001, ****p<0.0001, Two-way ANOVA with Šídák’s multiple comparisons test between preselected groups. **(B)** Relative expression of *per2, cry1a, per3,* and *per1b* in Z3 cells incubated with PBS or HK-Spn for 4 hours and NAC for 6 hours (2-hour pre-treatment + 4 hours treatment). *p<0.05, ****p<0.0001, ANOVA with Šídák’s multiple comparisons test between preselected groups.

After exposure to HK-Spn, ROS production was detected in all conditions (Fig.5A) and was highly augmented with increased light intensities. The kinetics of HK-Spn-induced ROS were light-dependent, such that faster kinetics were observed under high-intensity light than low-intensity light. Contrary to expectation, 500-lux light alone was not sufficient to induce ROS, indicating that ROS generated in response to HK-Spn results primarily from the detection of this microbe and not from the oxidative effects of light. In contrast, high-intensity light generated significant ROS after 3 hours, and adding HK-Spn further increased ROS. Light can directly react with tissue culture media components such as riboflavin to generate ROS, causing a false positive DCF signal^52^. We observed this effect in L15 media but not in HBSS (Fig.S2), and accordingly, these assays were all done in HBSS. As a control, the antioxidant N-acetylcysteine (NAC)^53^ was added to all conditions, and it significantly blocked the production of ROS induced by HK-Spn and high-intensity light (Fig 5). The fact that 500-lux light does not generate ROS (Fig. 5) but induces *per2* and *cry1a* (Fig. 1-3) suggests that ROS is not the primary driver of light-induced expression but may instead function as a secondary enhancer.

Based on this, we wondered whether HK-Spn-induced ROS contributes to the expression of circadian genes. To test this, we assessed NAC’s capacity to block HK-Spn effect on gene expression. Indeed, NAC effectively blocked HK-Spn’s ability to increase *per2*, *cry1a,* and *per3* expression at 4 hours, suggesting that ROS plays a role in this effect (Fig. 5B). NAC did not inhibit the induction of *per2* by light, supporting the idea that ROS is not the primary mechanism by which light induces *per2*. Additionally, this may explain why HK-Spn-mediated ROS induction has an additive effect with light but not with exogenous sources of ROS, such as H_2_O_2_. Unexpectedly, NAC increased the expression of *per1b* in cells exposed to light and HK-Spn (Fig. 5B). Additional studies showed the same effect of NAC on *per1b* in the dark (data not shown), suggesting this may result from unappreciated roles for antioxidants or reducing agents in the regulation of *per1b*.

### HK-Spn enhances the light-regulated expression of *tefa* and *tefb*

Another factor that has been implicated in the light-induction pathway is *tefa*^14^. This transcription factor is induced by light and triggers *per2* and *cry1a* expression by binding to D-boxes in their promoters^9,14,15,18,27,28^. The light-induction of *per2* is decreased when *tefb,* a homolog of *tefa*, is knocked-down, however it is unclear whether this is a direct effect on D-boxes^28,54^. Since the acute effects of HK Spn are enhanced in the presence of light, we wondered if these transcription factors participate in this process. We first examined whether the presence of bacteria impacts the expression of these genes. As expected, *tefa* was induced by light in an intensity-dependent manner (∼2-fold and ∼3.6-fold, respectively) (Fig.6A). Similar to the effect observed with *per2, cry1a,* and *per3,* the presence of HK-Spn augmented the light-induced expression of *tefa* at ∼500-lux (∼3.7-fold) and ∼5000-lux (∼7.2-fold). Surprisingly, light had a repressive effect on *tefb* which has not been previously reported in the literature. Intriguingly, HK-Spn induced expression of *tefb* in all conditions tested, including the dark and was even able to reverse the light-mediated repression of *tefb.* Given that HK-Spn can induce *tefa* and *tefb*, it is possible they may contribute to HK-Spn-mediated enhancement of *per2*, *cry1a,* and *per3*.

**Figure 6.**
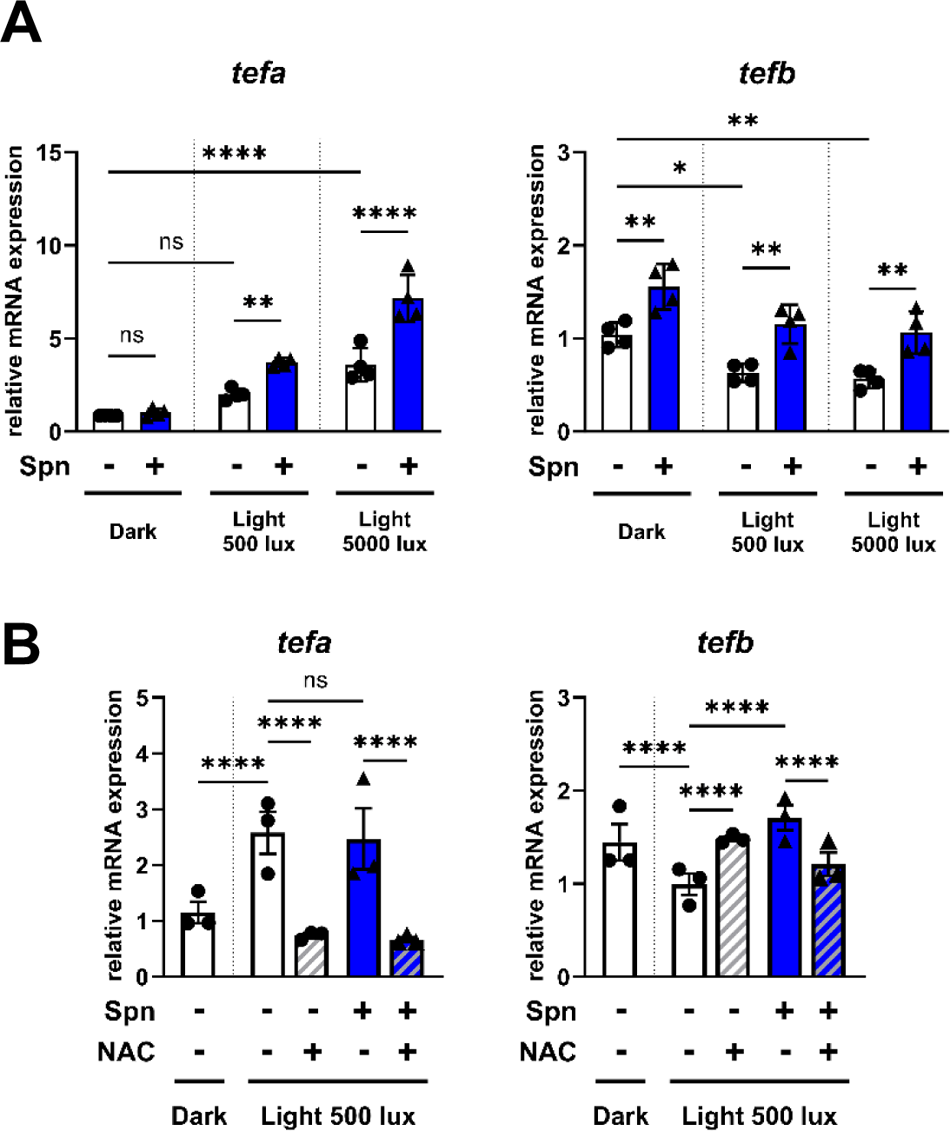
*tefa* and *tefb* are regulated by light, HK-Spn, and ROS. **(A)** Z3 cells were maintained in Z3 media in the dark, low-intensity light (∼500-lux) and high-intensity light (∼5000-lux) for 4 hours with PBS or HK-Spn. Relative RNA expression of *tefa* and *tefb* was measured after 4 hours. *p<0.05,** p<0.01, ****p<0.0001, Two-way ANOVA with Šídák’s multiple comparisons test of preselected groups. **(B)** Z3 cells were maintained in Z3 media in the dark or low-intensity light (∼500-lux) for 4 hours with PBS or HK-Spn. NAC was added as a 2-hour pre-treatment and continued after 4 hours of PBS/HK-Spn incubation. Relative RNA expression of *tefa* and *tefb* was measured after 4 hours. ****p<0.0001, Two-way ANOVA with Šídák’s multiple comparisons test of preselected

As we previously demonstrated the role of HK-Spn-induced ROS in augmenting *per2*, *cry1a*, and *per3*, we speculated that ROS might also contribute to HK-Spn-induced transcription of *tefa* and *tefb* and could be abrogated by NAC. As expected, *tefa* was induced by light after 4 hours (Fig.6B). NAC potently blocked light’s ability to induce *tefa* suggesting that ROS does play a role in the induction of *tefa.* In this experiment, unfortunately, we did not detect any augmentation in the expression of *tefa* caused by HK-Spn, possibly due to variation in bacteria preparations, which can induce a weaker or more potent effect on HK-Spn augmentation of genes. Despite this, the bacterial stock used was still sufficient to induce *tefb* with the same pattern seen in Figure 6A, and NAC was able to block this effect.

It is puzzling why NAC would have an effect on *tefa* or *tefb* when we didn’t detect any substantial fluorescence in DCF with these light conditions (Fig 5A). This suggests the presence of an ROS species generated by light that is not detectable by DCF. Moreover, it is possible that *tefb* is regulated by intricate redox pathways that trigger different expression outcomes depending on the ROS origin. For instance, light-induced ROS and HK-Spn-induced ROS may come from different sources (NOX, DUOX, etc.) or cell localization (cell membrane, mitochondria, nuclear membrane, etc.), resulting in the activation of various downstream pathways that will have different effects on *tefb* expression. Alternatively, NAC’s reducing activity can affect myriad biological molecules, any of which may affect gene expression. Nonetheless, these studies position microbial induced ROS in the regulation of core clock genes, perhaps through the actions of *tefa* and *tefb*.

## Discussion

Several studies have demonstrated changes in circadian gene expression during infection with bacteria, viruses, and parasites^35–40^. However, it is unclear whether this effect is caused by the inflammation resulting from disease or results directly from host detection of the microorganism itself. In this study, we investigated the direct impact of heat-killed *Streptococcus pneumoniae* on the expression of *per1b, per2*, *per3,* and *cry1a* in zebrafish Z3 cells. The expression patterns of these genes under basal conditions after 12-hour light exposure or continuous darkness (Fig.1) confirmed previous reports of *per2* and *cry1a* being rapidly induced by light, whereas *per1b* and *per3* showed no induction during light exposure^10,14,15,18,21^. The presence of bacteria potently augmented expression of *per1b, per2*, *per3,* and *cry1a* in a timing and condition-dependent manner (Fig.2). Furthermore, the analysis of the acute induction of these genes within 4 hours of HK-Spn exposure demonstrated consistent augmentation of *per2*, *cry1a,* and *per3* in the presence of light, while no increase was seen in the dark at this timepoint (Fig.3). In line with the fact that *per1b* is inhibited by light^21^, HK-Spn was unable to augment its expression in the light. Investigation into the mechanisms behind the HK-Spn-augmentation of gene expression demonstrated that HK-Spn induced ROS, which was further augmented in the presence of light (Fig.5A). We showed that the addition of the antioxidant NAC prevented HK-Spn from inducing *per2*, *cry1a*, and *per3* (Fig.5B) which positioned ROS as a central effector of this response. Finally, we found that HK-Spn was sufficient to induce the transcription factors *tefa* and *tefb* in a manner that could also be blocked by NAC, further highlighting potential roles of bacterially induced ROS in the expression of circadian genes (Fig.6).

The current understanding of how *per* and *cry* genes are regulated in zebrafish is thought to be through promoter elements called E-boxes and D-boxes. Circadian regulation is initiated by Clock:Bmal dimerization and binding to E-boxes in the promoters of clock-controlled genes, which include *per1b, per2*, *per3,* and *cry1a*^9,11,21,55^. Light-induced regulation is associated with the activation of D-boxes. However, while *cry1a* only needs the presence of one D-box, *per2* has been shown to require both E-boxes and D-boxes to be induced by light^14,15,17^. Moreover, reports of the involvement of light-generated ROS have also been related to the light-driven expression of *per2* and *cry1a,* though we were not able to replicate this finding in Z3 cells^10,16,17,34^. Possible reasons for these discrepancies may be the use of different cell types (Z3 cells vs. PAC-2 cells), different light spectrums (white light vs. blue light) and intensities, or the fact that our assays were conducted in HBSS and therefore do not include ROS generated from light-oxidation of common tissue culture media components (Fig.S2)^52^. Regardless, the induction of *per2* and *cry1a* by 500-lux light without generating ROS implies that ROS are not the primary factor for the light-dependent transcription of these genes. This is supported by the fact that NAC did not block the light-induced expression of *per2*. However, we did see a blocking effect of NAC in *cry1a* light-induction, suggesting that light may generate a type of ROS that is quenched by NAC but is not detectable with DCF. For example, DCF can only detect cellular peroxides, such as H_2_O_2_, if they are decomposed to radicals; however, exogenous H_2_O_2_ can induce *per2* and *cry1a* (Fig.4)^56^. Furthermore, it is possible that the ROS generated by light and those induced by HK-Spn are not the same type since DCF can be oxidized by various radicalized ROS, including peroxyl, alkoxyl, hydroxyl, carbonate, nitrogen dioxide, and peroxynitrite^56^.

A well-known effect of pathogen recognition by innate immune cells is the production of ROS as an antimicrobial agent^32,33,50^. Here, we demonstrate that Z3 cells, a non-immune cell type, can also generate ROS in response to pathogen exposure. Interestingly, high-intensity light further increased HK-Spn-ROS production and correlated with even stronger expression of *per2*, *cry1a,* and *per3.* This, and the fact that NAC could block these phenotypes, strongly positions HK-Spn-induced ROS as an enhancer of expression and reaffirms the correlation between redox interactions and the circadian clock^16,17,57^. However, it is uncertain how ROS performs these changes. One possibility is that HK-Spn-induced ROS directly affects the function of proteins such as transcription factors by modification of cysteine residues susceptible to oxidation^32^. Alternatively, ROS may intersect with central signaling pathways responsible for activating transcription factors that target circadian genes. For instance, H_2_O_2_ has been linked to the activation of the MAP kinases p38 and JNK and D-boxes in PAC-2 cells and has been shown to increase levels of CLOCK, BMAL1, PER1/2, AND CRY1/2 through activation of the PRX2-STAT3-REV-ERBα/β pathway in NIH3T3 cells^17,58^

In searching for candidates that could participate in this regulation, we performed a comparative analysis of putative binding sites of different transcription factors in the promoter regions of *per1b, per2*, *per3,* and *cry1a* (not shown). Binding sites for circadian transcription factors (D-box, E-box), as well as sites for a variety of non-circadian-related transcription factors (such as NF-kB, AP-1, HIF-1α, etc.), were identified in the promoters of these genes, but unfortunately, we were unable to detect any obvious pattern to explain the observed results. Despite this, the strongest phenotype observed in our data occurs in a condition when D-boxes are active: *per2* and *cry1a* induction in the light. Thus, we focused our investigation on the transcription factors *tefa* and *tefb*, which have been implicated in circadian modulation and light-regulation of *per2* and *cry1a*^14,27,28,54^. Previously, it has been shown that *tefa* is induced by light while *tefb* troughs during the light period^28^. Here, we demonstrate that *tefb* is actually repressed by light. Our results indicate that HK-Spn can augment light-induced *tefa* and induce *tefb* even under the repressing effect of light. It remains unclear whether HK-Spn induced expression of *tefa* is sufficient to enhance *per2* and *cry1a*, or whether *tefb* may contribute to this phenotype.

It is currently unknown whether *tefb* can bind to D-boxes, however evidence of the participation of both *tefa* and *tefb* in the light regulation of *per2* is supported by studies showing that morpholino knockdown either of *tefa* and *tefb* decreased the light-induced expression of *per2*^28^. *Tefa* is a light-induced gene which can drive expression of *per2* via D-boxes, an effect that is additive with Clock:Bmal mediated expression via E-boxes^14,18^. Since *per2* and *cry1a* have D-boxes, the HK-Spn-induced augmentation of *tefa* alone could explain the increase in these genes. However, this does not explain how *per3* is augmented as it does not possess D-boxes. It is possible that other transcription factors are responsible, such as NF-kB, AP-1, HIF-1α, or *tefb*, who’s binding sites have not been identified. Additionally, other factors such as promoter architecture, i.e., combination and spacing of binding sites or interactions between factors, could be at play in this regulation^55,59,60^. Regardless, the findings presented in this work provide new insights into the regulatory pathways of zebrafish circadian genes and present, for the first time, a possible mechanism of microbe-circadian clock interaction in zebrafish cells through ROS. Understanding the interactions between different types of bacteria and the circadian clock can provide valuable insights into the intricate relationship between these systems.

## Acknowledgements

This work was supported by the National Institutes of Health / NIGMS (R35GM147509). J.M.K. is a Pew Scholar in the Biomedical Sciences, supported by The Pew Charitable Trusts.

---

**Figure S1.**
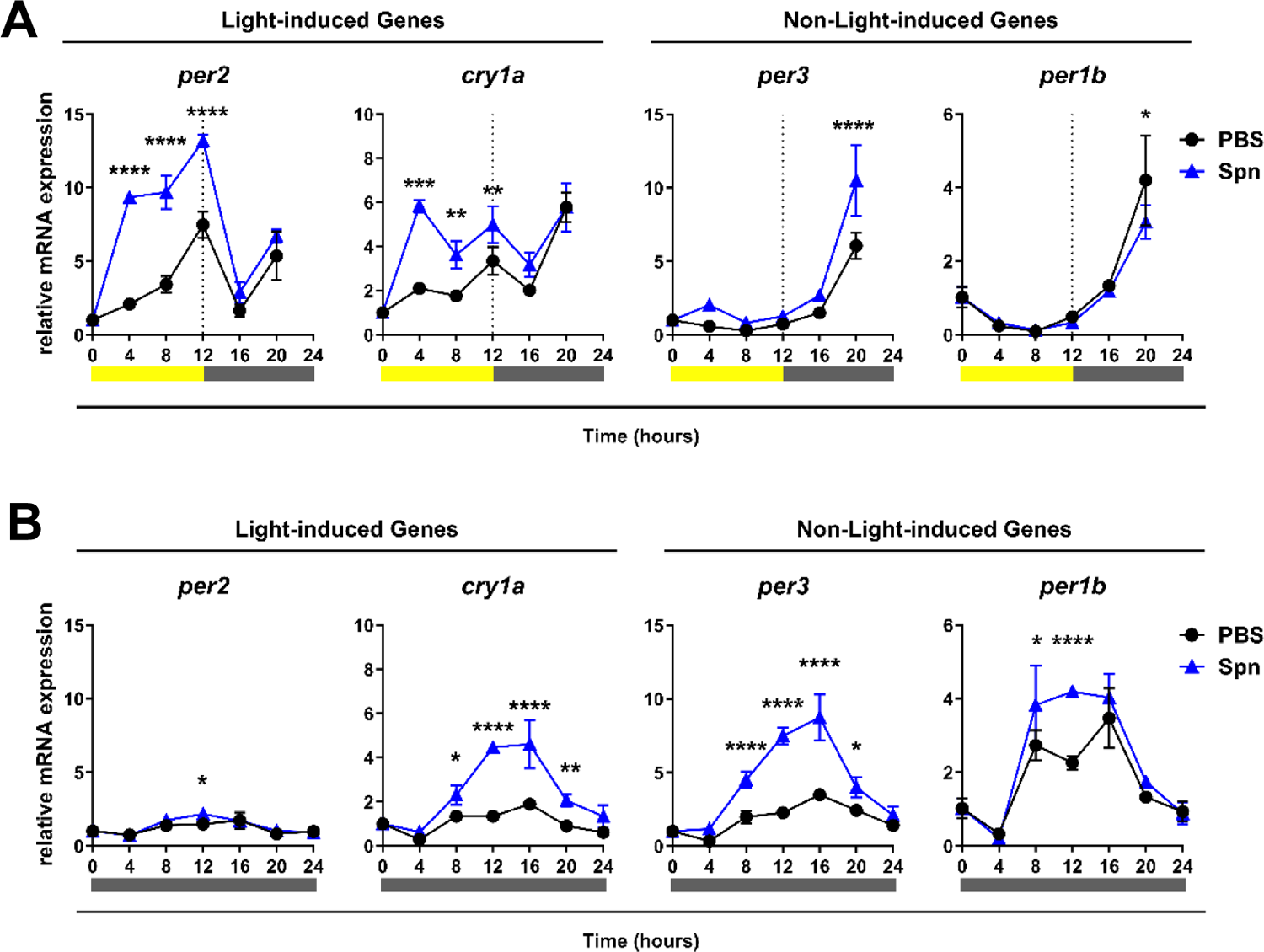
Expression of *per2, cry1a, per3* and *per1b* at the first 24 hours of HK HK-Spn exposure. Z3 cells were entrained for three LD cycles of 12:12 h, followed by a 12-hour light pulse (LDDD) or in constant darkness (DDDD). Heat-killed *Streptococcus pneumoniae* (HK HK-Spn) was added at time 0, and samples were collected every 4 hours throughout 48 hours. **(A)** Transcript expression of *per2, cry1a*, *per3*, and *per1b* after 12-hour light (500-lux) followed by constant darkness of PBS or HK HK-Spn exposed cells. **(B)** Transcript expression of *per2, cry1a*, *per3,* and *per1b* in constant darkness of PBS or HK HK-Spn exposed cells. *p<0.05,** p<0.01,***p<0.001, ****p<0.0001, Two-way ANOVA with Šídák’s multiple comparisons test. Yellow bar: light period. Grey bar: dark period.

**Figure S2.**
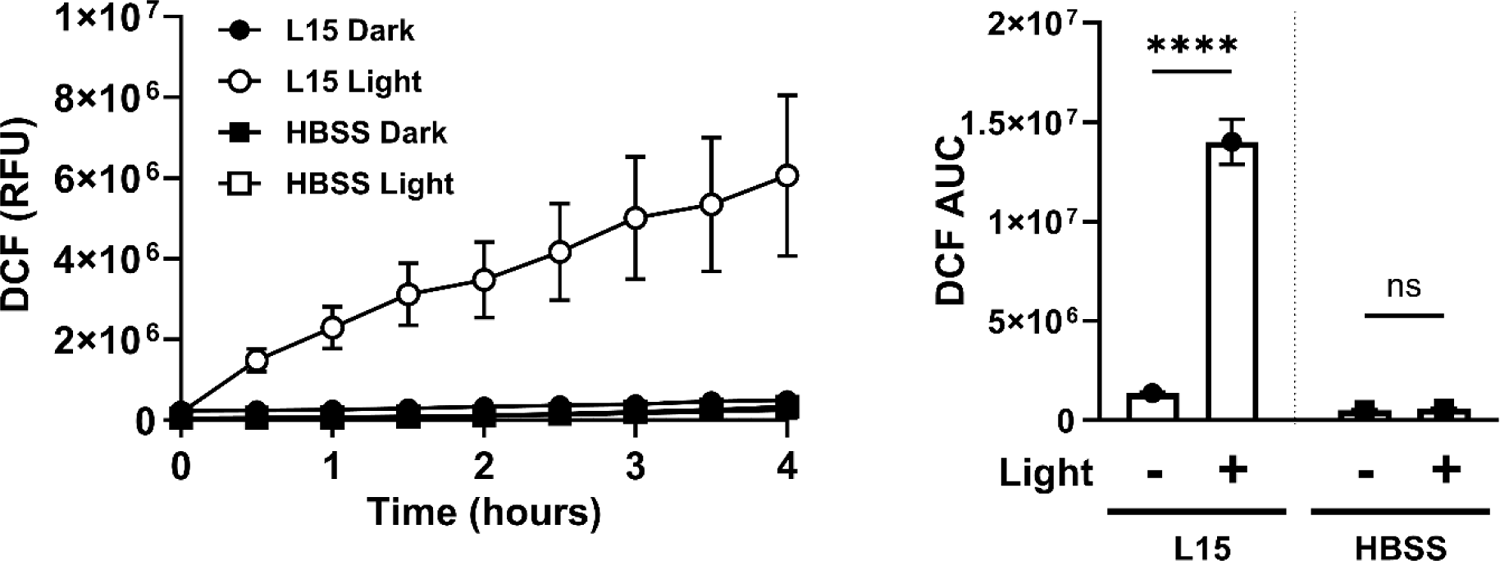
Light generates ROS in L15 media alone but not in HBSS. Relative fluorescent units (RFU) of Dichlorohydrofluoresin (DCF) were measured every 30 minutes for 4 hours on L15 media Hank’s or Balanced Salt Solution (HBSS), in the absence of cells, in the dark and in low-intensity light (∼500-lux). **(C)** The area under the curve (AUC) was measured and compared between groups and conditions. *p<0.05, ***p<0.001, ****p<0.0001, Two-way ANOVA with Šídák’s multiple comparisons test between preselected groups.

## References

(1) Partch, C. L.; Green, C. B.; Takahashi, J. S. Molecular Architecture of the Mammalian Circadian Clock. Trends in Cell Biology 2014, 24 (2), 90–99. 10.1016/j.tcb.2013.07.002.

(2) Boivin, D. B.; Duffy, J. F.; Kronauer, R. E.; Czeisler, C. A. Dose-Response Relationships for Resetting of Human Circadian Clock by Light. Nature 1996, 379 (6565), 540–542. 10.1038/379540a0.

(3) Hannay, K. M.; Forger, D. B.; Booth, V. Seasonality and Light Phase-Resetting in the Mammalian Circadian Rhythm. Sci Rep 2020, 10 (1), 19506. 10.1038/s41598-020-74002-2.

(4) Jiménez, A.; Lu, Y.; Jambhekar, A.; Lahav, G. Principles, Mechanisms and Functions of Entrainment in Biological Oscillators. Interface Focus. 2022, 12 (3), 20210088. 10.1098/rsfs.2021.0088.

(5) Whitmore, D.; Foulkes, N. S.; Sassone-Corsi, P. Light Acts Directly on Organs and Cells in Culture to Set the Vertebrate Circadian Clock. Nature 2000, 404 (6773), 87–91. 10.1038/35003589.

(6) Kobayashi, Y.; Ishikawa, T.; Hirayama, J.; Daiyasu, H.; Kanai, S.; Toh, H.; Fukuda, I.; Tsujimura, T.; Terada, N.; Kamei, Y.; Yuba, S.; Iwai, S.; Todo, T. Molecular Analysis of Zebrafish Photolyase/Cryptochrome Family: Two Types of Cryptochromes Present in Zebrafish. Genes to Cells 2000, 5 (9), 725–738. 10.1046/j.1365-2443.2000.00364.x.

(7) Liu, C.; Hu, J.; Qu, C.; Wang, L.; Huang, G.; Niu, P.; Zhong, Z.; Hong, F.; Wang, G.; Postlethwait, J. H.; Wang, H. Molecular Evolution and Functional Divergence of Zebrafish (Danio Rerio) Cryptochrome Genes. Sci Rep 2015, 5 (1), 8113. 10.1038/srep08113.

(8) West, A. C.; Iversen, M.; Jørgensen, E. H.; Sandve, S. R.; Hazlerigg, D. G.; Wood, S. H. Diversified Regulation of Circadian Clock Gene Expression Following Whole Genome Duplication. PLoS Genet 2020, 16 (10), e1009097. 10.1371/journal.pgen.1009097.

(9) Vatine, G.; Vallone, D.; Gothilf, Y.; Foulkes, N. S. It’s Time to Swim! Zebrafish and the Circadian Clock. FEBS Letters 2011, 585 (10), 1485–1494. 10.1016/j.febslet.2011.04.007.

(10) Junko, I.; Okamoto-Uchida, Y.; Nishimura, A.; Hirayama, J. Light-Dependent Regulation of Circadian Clocks in Vertebrates. In *Chronobiology - The Science of Biological Time Structure*; Svorc, P., Ed.; IntechOpen, 2019. 10.5772/intechopen.86524.

(11) Sacksteder, R. E.; Kimmey, J. M. Immunity, Infection, and the Zebrafish Clock. Infect Immun 2022, 90 (9), e00588–21. 10.1128/iai.00588-21.

(12) Pando, M. P.; Pinchak, A. B.; Cermakian, N.; Sassone-Corsi, P. A Cell-Based System That Recapitulates the Dynamic Light-Dependent Regulation of the Vertebrate Clock. Proc Natl Acad Sci U S A 2001, 98 (18), 10178–10183. 10.1073/pnas.181228598.

(13) Tamai, T. K.; Young, L. C.; Whitmore, D. Light Signaling to the Zebrafish Circadian Clock by *Cryptochrome 1a*. Proc. Natl. Acad. Sci. U.S.A. 2007, 104 (37), 14712–14717. 10.1073/pnas.0704588104.

(14) Vatine, G.; Vallone, D.; Appelbaum, L.; Mracek, P.; Ben-Moshe, Z.; Lahiri, K.; Gothilf, Y.; Foulkes, N. S. Light Directs Zebrafish Period2 Expression via Conserved D and E Boxes. PLoS Biol 2009, 7 (10), e1000223. 10.1371/journal.pbio.1000223.

(15) Mracek, P.; Santoriello, C.; Idda, M. L.; Pagano, C.; Ben-Moshe, Z.; Gothilf, Y.; Vallone, D.; Foulkes, N. S. Regulation of per and Cry Genes Reveals a Central Role for the D-Box Enhancer in Light-Dependent Gene Expression. PLOS ONE 2012, 7 (12), e51278. 10.1371/journal.pone.0051278.

(16) Hirayama, J.; Cho, S.; Sassone-Corsi, P. Circadian Control by the Reduction/Oxidation Pathway: Catalase Represses Light-Dependent Clock Gene Expression in the Zebrafish. PNAS 2007, 104 (40), 15747–15752.

(17) Pagano, C.; Siauciunaite, R.; Idda, M. L.; Ruggiero, G.; Ceinos, R. M.; Pagano, M.; Frigato, E.; Bertolucci, C.; Foulkes, N. S.; Vallone, D. Evolution Shapes the Responsiveness of the D-Box Enhancer Element to Light and Reactive Oxygen Species in Vertebrates. Sci Rep 2018, 8 (1), 13180. 10.1038/s41598-018-31570-8.

(18) Weger, B. D.; Sahinbas, M.; Otto, G. W.; Mracek, P.; Armant, O.; Dolle, D.; Lahiri, K.; Vallone, D.; Ettwiller, L.; Geisler, R.; Foulkes, N. S.; Dickmeis, T. The Light Responsive Transcriptome of the Zebrafish: Function and Regulation. PLoS ONE 2011, 6 (2), e17080. 10.1371/journal.pone.0017080.

(19) Moore, H. A.; Whitmore, D. Circadian Rhythmicity and Light Sensitivity of the Zebrafish Brain. PLoS ONE 2014, 9 (1), e86176. 10.1371/journal.pone.0086176.

(20) Albrecht, U.; Zheng, B.; Larkin, D.; Sun, Z. S.; Lee, C. C. *mPer1* and *mPer2* Are Essential for Normal Resetting of the Circadian Clock. J Biol Rhythms 2001, 16 (2), 100–104. 10.1177/074873001129001791.

(21) Vallone, D.; Gondi, S. B.; Whitmore, D.; Foulkes, N. S. E-Box Function in a Period Gene Repressed by Light. PNAS 2004, 101 (12), 4106–4111. 10.1073/pnas.0305436101.

(22) Frøland Steindal, I.; Whitmore, D. Circadian Clocks in Fish—What Have We Learned so Far? Biology 2019, 8 (1), 17. 10.3390/biology8010017.

(23) Moutsaki, P.; Whitmore, D.; Bellingham, J.; Sakamoto, K.; David-Gray, Z. K.; Foster, R. G. Teleost Multiple Tissue (Tmt) Opsin: A Candidate Photopigment Regulating the Peripheral Clocks of Zebrafish? Molecular Brain Research 2003, 112 (1–2), 135–145. 10.1016/S0169-328X(03)00059-7.

(24) Cavallari, N.; Frigato, E.; Vallone, D.; Fröhlich, N.; Lopez-Olmeda, J. F.; Foà, A.; Berti, R.; Sánchez-Vázquez, F. J.; Bertolucci, C.; Foulkes, N. S. A Blind Circadian Clock in Cavefish Reveals That Opsins Mediate Peripheral Clock Photoreception. PLOS Biology 2011, 9 (9), e1001142. 10.1371/journal.pbio.1001142.

(25) Ozturk, N. Light-dependent Reactions of Animal Circadian Photoreceptor Cryptochrome. The FEBS Journal 2022, 289 (21), 6622–6639. 10.1111/febs.16273.

(26) Gachon, F. Physiological Function of PARbZip Circadian Clock-controlled Transcription Factors. Annals of Medicine 2007, 39 (8), 562–571. 10.1080/07853890701491034.

(27) Ben-Moshe, Z.; Vatine, G.; Alon, S.; Tovin, A.; Mracek, P.; Foulkes, N. S.; Gothilf, Y. MULTIPLE PAR AND E4BP4 bZIP TRANSCRIPTION FACTORS IN ZEBRAFISH: DIVERSE SPATIAL AND TEMPORAL EXPRESSION PATTERNS. Chronobiology International 2010, 27 (8), 1509–1531. 10.3109/07420528.2010.510229.

(28) Gavriouchkina, D.; Fischer, S.; Ivacevic, T.; Stolte, J.; Benes, V.; Dekens, M. P. S. Thyrotroph Embryonic Factor Regulates Light-Induced Transcription of Repair Genes in Zebrafish Embryonic Cells. PLoS ONE 2010, 5 (9), e12542. 10.1371/journal.pone.0012542.

(29) Ichikawa, M.; Nishino, T.; Nishino, T.; Ichikawa, A. Subcellular Localization of Xanthine Oxidase in Rat Hepatocytes: High-Resolution Immunoelectron Microscopic Study Combined with Biochemical Analysis. J Histochem Cytochem. 1992, 40 (8), 1097–1103. 10.1177/40.8.1619276.

(30) Donkó, Á.; Péterfi, Z.; Sum, A.; Leto, T.; Geiszt, M. Dual Oxidases. Phil. Trans. R. Soc. B 2005, 360 (1464), 2301–2308. 10.1098/rstb.2005.1767.

(31) Skonieczna, M.; Hejmo, T.; Poterala-Hejmo, A.; Cieslar-Pobuda, A.; Buldak, R. J. NADPH Oxidases: Insights into Selected Functions and Mechanisms of Action in Cancer and Stem Cells. Oxidative Medicine and Cellular Longevity 2017, 2017, 1–15. 10.1155/2017/9420539.

(32) Sies, H.; Jones, D. P. Reactive Oxygen Species (ROS) as Pleiotropic Physiological Signalling Agents. Nat Rev Mol Cell Biol 2020, 21 (7), 363–383. 10.1038/s41580-020-0230-3.

(33) Li, P.; Chang, M. Roles of PRR-Mediated Signaling Pathways in the Regulation of Oxidative Stress and Inflammatory Diseases. IJMS 2021, 22 (14), 7688. 10.3390/ijms22147688.

(34) Uchida, Y.; Hirayama, J.; Nishina, H. A Common Origin: Signaling Similarities in the Regulation of the Circadian Clock and DNA Damage Responses. Biological & Pharmaceutical Bulletin 2010, 33 (4), 535–544. 10.1248/bpb.33.535.

(35) Benegiamo, G.; Mazzoccoli, G.; Cappello, F.; Rappa, F.; Scibetta, N.; Oben, J.; Greco, A.; Williams, R.; Andriulli, A.; Vinciguerra, M.; Pazienza, V. Mutual Antagonism between Circadian Protein Period 2 and Hepatitis C Virus Replication in Hepatocytes. PLoS ONE 2013, 8 (4), e60527. 10.1371/journal.pone.0060527.

(36) Li, T.; Shao, W.; Li, S.; Ma, L.; Zheng, L.; Shang, W.; Jia, X.; Sun, P.; Liang, X.; Jia, J. H. Pylori Infection Induced BMAL1 Expression and Rhythm Disorder Aggravate Gastric Inflammation. EBioMedicine 2019, 39, 301–314. 10.1016/j.ebiom.2018.11.043.

(37) Huang, H.; Mehta, A.; Kalmanovich, J.; Anand, A.; Bejarano, M. C.; Garg, T.; Khan, N.; Tonpouwo, G. K.; Shkodina, A. D.; Bardhan, M. Immunological and Inflammatory Effects of Infectious Diseases in Circadian Rhythm Disruption and Future Therapeutic Directions. Mol Biol Rep 2023, 50 (4), 3739– 3753. 10.1007/s11033-023-08276-w.

(38) Rijo-Ferreira, F.; Carvalho, T.; Afonso, C.; Sanches-Vaz, M.; Costa, R. M.; Figueiredo, L. M.; Takahashi, J. S. Sleeping Sickness Is a Circadian Disorder. Nat Commun 2018, 9 (1), 62. 10.1038/s41467-017-02484-2.

(39) Ellison, A. R.; Wilcockson, D.; Cable, J. Circadian Dynamics of the Teleost Skin Immune-Microbiome Interface. Microbiome 2021, 9 (1), 222. 10.1186/s40168-021-01160-4.

(40) Midttun, H. L. E.; Vindas, M. A.; Whatmore, P. J.; Øverli, Ø.; Johansen, I. B. Effects of *Pseudoloma Neurophilia* Infection on the Brain Transcriptome in Zebrafish ( *Danio Rerio*). Journal of Fish Diseases 2020, 43 (8), 863–875. 10.1111/jfd.13198.

(41) Chen, S.; Fuller, K. K.; Dunlap, J. C.; Loros, J. J. A Pro- and Anti-Inflammatory Axis Modulates the Macrophage Circadian Clock. Front. Immunol. 2020, 11, 867. 10.3389/fimmu.2020.00867.

(42) Cavadini, G.; Petrzilka, S.; Kohler, P.; Jud, C.; Tobler, I.; Birchler, T.; Fontana, A. TNF-α Suppresses the Expression of Clock Genes by Interfering with E-Box-Mediated Transcription. Proc. Natl. Acad. Sci. U.S.A. 2007, 104 (31), 12843–12848. 10.1073/pnas.0701466104.

(43) Kwak, Y.; Lundkvist, G. B.; Brask, J.; Davidson, A.; Menaker, M.; Kristensson, K.; Block, G. D. Interferon-γ Alters Electrical Activity and Clock Gene Expression in Suprachiasmatic Nucleus Neurons. J Biol Rhythms 2008, 23 (2), 150–159. 10.1177/0748730407313355.

(44) Wang, Y.; Pati, P.; Xu, Y.; Chen, F.; Stepp, D. W.; Huo, Y.; Rudic, R. D.; Fulton, D. J. R. Endotoxin Disrupts Circadian Rhythms in Macrophages via Reactive Oxygen Species. PLoS ONE 2016, 11 (5), e0155075. 10.1371/journal.pone.0155075.

(45) Mosser, E. A.; Chiu, C. N.; Tamai, T. K.; Hirota, T.; Li, S.; Hui, M.; Wang, A.; Singh, C.; Giovanni, A.; Kay, S. A.; Prober, D. A. Identification of Pathways That Regulate Circadian Rhythms Using a Larval Zebrafish Small Molecule Screen. Sci Rep 2019, 9. 10.1038/s41598-019-48914-7.

(46) Livak, K. J.; Schmittgen, T. D. Analysis of Relative Gene Expression Data Using Real-Time Quantitative PCR and the 2−ΔΔCT Method. Methods 2001, 25 (4), 402–408. 10.1006/meth.2001.1262.

(47) Delaunay, F.; Thisse, C.; Marchand, O.; Laudet, V.; Thisse, B. An Inherited Functional Circadian Clock in Zebrafish Embryos. Science 2000, 289 (5477), 297–300. 10.1126/science.289.5477.297.

(48) Kaneko, M.; Cahill, G. M. Light-Dependent Development of Circadian Gene Expression in Transgenic Zebrafish. PLoS Biol 2005, 3 (2), e34. 10.1371/journal.pbio.0030034.

(49) Yang, Y.; Bazhin, A. V.; Werner, J.; Karakhanova, S. Reactive Oxygen Species in the Immune System. International Reviews of Immunology 2013, 32 (3), 249–270. 10.3109/08830185.2012.755176.

(50) Martinvalet, D.; Walch, M. Editorial: The Role of Reactive Oxygen Species in Protective Immunity. Front. Immunol. 2022, 12, 832946. 10.3389/fimmu.2021.832946.

(51) Gardiner, B.; Dougherty, J. A.; Ponnalagu, D.; Singh, H.; Angelos, M.; Chen, C.-A.; Khan, M. Measurement of Oxidative Stress Markers In Vitro Using Commercially Available Kits. In Measuring Oxidants and Oxidative Stress in Biological Systems; Berliner, L. J., Parinandi, N. L., Eds.; Biological Magnetic Resonance; Springer International Publishing: Cham, 2020; Vol. 34, pp 39–60. 10.1007/978-3-030-47318-1_4.

(52) Grzelak, A.; Rychlik, B.; Bartosz, G. Light-Dependent Generation of Reactive Oxygen Species in Cell Culture Media. Free Radical Biology and Medicine 2001, 30 (12), 1418–1425. 10.1016/S0891-5849(01)00545-7.

(53) Sun, S.-Y. N-Acetylcysteine, Reactive Oxygen Species and Beyond. Cancer Biology & Therapy 2010, 9 (2), 109–110. 10.4161/cbt.9.2.10583.

(54) Gachon, F.; Olela, F. F.; Schaad, O.; Descombes, P.; Schibler, U. The Circadian PAR-Domain Basic Leucine Zipper Transcription Factors DBP, TEF, and HLF Modulate Basal and Inducible Xenobiotic Detoxification. Cell Metabolism 2006, 4 (1), 25–36. 10.1016/j.cmet.2006.04.015.

(55) Gordân, R.; Shen, N.; Dror, I.; Zhou, T.; Horton, J.; Rohs, R.; Bulyk, M. L. Genomic Regions Flanking E-Box Binding Sites Influence DNA Binding Specificity of bHLH Transcription Factors through DNA Shape. Cell Reports 2013, 3 (4), 1093–1104. 10.1016/j.celrep.2013.03.014.

(56) Eruslanov, E.; Kusmartsev, S. Identification of ROS Using Oxidized DCFDA and Flow-Cytometry. In Advanced Protocols in Oxidative Stress II; Armstrong, D., Ed.; Methods in Molecular Biology; Humana Press: Totowa, NJ, 2010; Vol. 594, pp 57–72. 10.1007/978-1-60761-411-1_4.

(57) Lai, A. G.; Doherty, C. J.; Mueller-Roeber, B.; Kay, S. A.; Schippers, J. H. M.; Dijkwel, P. P. *CIRCADIAN CLOCK-ASSOCIATED 1* Regulates ROS Homeostasis and Oxidative Stress Responses. Proc. Natl. Acad. Sci. U.S.A. 2012, 109 (42), 17129–17134. 10.1073/pnas.1209148109.

(58) Ji, G.; Lv, K.; Chen, H.; Wang, Y.; Zhang, Y.; Li, Y.; Qu, L. Hydrogen Peroxide Modulates Clock Gene Expression via PRX2-STAT3-REV-ERBα/β Pathway. Free Radical Biology and Medicine 2019, 145, 312–320. 10.1016/j.freeradbiomed.2019.09.036.

(59) Nakahata, Y.; Yoshida, M.; Takano, A.; Soma, H.; Yamamoto, T.; Yasuda, A.; Nakatsu, T.; Takumi, T. A Direct Repeat of E-Box-like Elements Is Required for Cell-Autonomous Circadian Rhythm of Clock Genes. BMC Molecular Biol 2008, 9 (1), 1. 10.1186/1471-2199-9-1.

(60) Cox, K. H.; Takahashi, J. S. Circadian Clock Genes and the Transcriptional Architecture of the Clock Mechanism. Journal of Molecular Endocrinology 2019, 63 (4), R93–R102. 10.1530/JME-19-0153.

